# Unbiased recording of clonal potency reveals species-specific regulation of mammalian intestine

**DOI:** 10.1101/2025.06.10.658707

**Authors:** Mirazul Islam, Matthew E. Bechard, Yilin Yang, Alan J. Simmons, Yanwen Xu, James N. Higginbotham, Ping Zhao, Zheng Cao, Naila Tasneem, Sarah E. Glass, Nicholas O. Markham, Frank Revetta, Marisol A. Ramirez-Solano, Qi Liu, Jeffrey L. Franklin, Ken S. Lau, Robert J. Coffey

## Abstract

The mammalian intestine regenerates rapidly after damage, yet the clonal dynamics and species-specific regulation of different populations remain poorly understood. Here we used synthetic or naturally occurring DNA alterations to reconstruct clonal histories of the mouse and human intestinal epithelium at single-cell resolution. In mice, we uncovered the clonal architecture of different cell types and their roles in regeneration, supporting a hierarchical regenerative response model. We identified a rare embryonic precursor population that persisted in the adult and was crucial for regeneration after irradiation. This population was marked by Tob2, which is required for nuclear transport of Ascl2. A parallel clonal analysis of 65 human colonic biopsies revealed secretory lineage bias and an age-associated decline in clonal diversity in the distal colon. Unlike highly proliferative murine Lgr5+ stem cells, human LGR5+ cells were found largely quiescent, revealing species-specific difference in clonal potency, and suggesting a distinct regulation of intestinal stemness.

## INTRODUCTION

Based on studies in the mouse, rapidly proliferative Lgr5^+^ crypt-base columnar cells (CBCs) are widely recognized as the homeostatic intestinal stem cell (ISC) with the defining properties of continuous self-renewal and multipotency [1]. However, it is unclear whether human LGR5^+^ cells are equivalent to murine Lgr5^+^ ISCs, given the inability of performing *in vivo* lineage labelling in humans. Moreover, recent findings in mouse models challenge the assumption that adult CBCs are exclusively derived from embryonic Lgr5^+^ precursors [2], implying that other embryonic populations are precursors to the adult stem cell pool. Although CBCs are fast-cycling and sensitive to insults, such as irradiation (IR) [3, 4], the intestinal epithelium still rapidly regenerates following damage [5, 6]. Intriguingly, Lgr5^+^ cells are dispensable in homeostasis [7], yet required for radiation-induced intestinal regeneration [6], suggesting a dynamic interplay between different cell populations during injury repair. Multiple cell populations have been reported to contribute to epithelial regeneration after damage [8-10], thereby maintaining epithelial barrier function. How different regenerative populations accomplish both the replenishment of CBCs and the production of differentiated cells—two seemingly opposing tasks—within a short time frame remains unclear. Evidence supports the co-existence of quiescent and active stem cells within the crypt [11]; however, the precise mechanisms regulating quiescence in a high-turnover tissue system like the intestine remain poorly understood. The dynamics between quiescent and active states and their contribution to cells within the intestinal epithelium likely plays a critical role in balancing epithelial maintenance and regeneration, particularly under conditions of stress or injury.

The traditional experimental approach to assess stem cell contribution to tissues is prospective lineage tracing, such as the Cre recombinase approach where progenies arising from a candidate stem cell population are genetically marked and visualized [12, 13]. A general caveat to this approach is the dependence on a pre-selected promoter to trace from a single cell type, where the differential contributions originating from multiple cell populations—a new measure we call clonal potency— cannot be quantitatively compared. This, along with other technical variables such as non-specific promoter activity and variable Cre recombination efficiencies, limit the ability to explore unbiased clonal dynamics during homeostasis and regeneration [14-17]. Furthermore, while genetic labeling can be performed in human intestinal organoids or xenografts [18, 19], such exogenous labels cannot be applied to human tissues *in vivo*. As a result, our understanding of the quantitative contributions of different cell types and states for high-turnover tissues, such as the intestine, remains limited. Here, we define clonal potency as a cell-intrinsic property that quantifies the contribution of that cell’s progeny to the differentiated tissue.

Recent studies have shown that spontaneously acquired or induced somatic mutations act as indelible genetic barcodes, enabling retrospective, unbiased lineage tracing in both humans and mice [20-27]. Our group, and others, have previously demonstrated that CRISPR-induced mutations are passed on to daughter cells after division and they accumulate over mitotic cycles, allowing for the tracking of clonal events and cell lineages throughout murine development [21, 22, 28]. While this approach has been applied in various tissues, it has yet to be used for exploring the mammalian intestinal epithelium at single-cell resolution. Here, we leverage this strategy [28] to investigate cellular origins and clonal dynamics of the rapidly renewing intestinal epithelium during homeostasis and following IR-induced regeneration. Through this approach, we identified and characterized a distinct cell population called persister intestinal stem cells (pISCs), which are specified during early embryogenesis and persist in the adult intestine and after IR, playing a critical role in regeneration. Using human mitochondrial mutational phylogenies paired with transcriptional states, our analysis of the normal human colonic epithelium uncovered the quiescent nature of human LGR5^+^ cells, along with region-specific lineage biases and age-associated shifts in clonal diversity *in vivo*.

## RESULTS

### Clonal potency from quiescent and cycling cells in intestinal regeneration

Fast-cycling cells with high Wnt activity die after IR [3], and the remaining survivor cells replenish the damaged epithelium. We considered two possibilities that could occur during regeneration: (i) survivor cells regenerate CBCs, which then give rise to differentiated cells, or (ii) survivor cells generate both CBCs and differentiated cells concurrently (Figure 1A). To distinguish between these two possibilities, we administered whole-body IR to developmentally barcoded mice [20] and profiled the regenerated epithelial population using native single-guide RNA capture and sequencing (NSC-seq) [28], which captures lineage barcodes and transcriptomes simultaneously (Figures 1B, S1A, and S2B). We identified known intestinal epithelial cell types alongside pISCs (Figures 1C, 1D, and S1C-S1F), which exhibit a unique genetic phylogeny, suggesting their origin in early embryonic development and persistence in adult tissue [28]. A subset of the pISC population is found to be more proliferative after IR (pISC2s), while pISC1s remain quiescent. Clonal analysis performed on all epithelial cells revealed a significant increase in clone size and proliferation, accompanied by a decrease in clonal diversity after IR, reflecting the expected cellular loss following IR and clonal expansion during regeneration (Figure S1G). We identified pISC2 as a cell population that increases in absolute number, as well as clonal fraction after IR (Figures 1E and 1F). The distribution of pISC2-rooted clones increased three-fold, along with a rise in the pISC2 parent clone proportion after IR, implying an enhanced contribution to regeneration (Figures 1G-1I and S1H). Within the same animals, we also examined the clonal dynamics of CBCs in regeneration. If CBCs are obligatory intermediate cells during regeneration, we would expect to see an increased representation of CBC-rooted clones across differentiated cell types. However, to our surprise, we found the CBC-rooted clones decrease after regeneration (Figures 1J, 1K, and S1I), suggesting that CBCs are not an obligatory intermediate progenitor during epithelial regeneration. Analysis of clonal potency of individual cell types after IR revealed four populations (TA, pISC2, Paneth cells, and pISC1) actively contributing to regeneration (Figure 1L). Among them, Paneth cells have been shown to contribute to IR-induced regeneration [8]. A comparative analysis of the reconstructed single-cell lineage trees suggested clonal expansion after IR, where both Paneth cells and pISC2 independently contribute to regeneration (Figures S1J-S1L). Finally, we observed that pISC2 showed heightened Wnt and Notch signaling after IR, along with increased proliferation [8]. pISC2 shared a phylogenetic relationship with CBCs, suggesting a replenishment of CBCs by pISC2 following IR, underscoring its contribution to regeneration (Figures S1M-S1O). Together, these data provide mechanistic insights into the interplay between cell populations that contribute to IR-induced regeneration and suggest pISC2 is a major driver of intestinal regeneration (Figure S1P). Taken together, these results support the second possibility proposed, that is, that survivor cells generate CBCs and differentiated cells concurrently.

**Figure 1.**
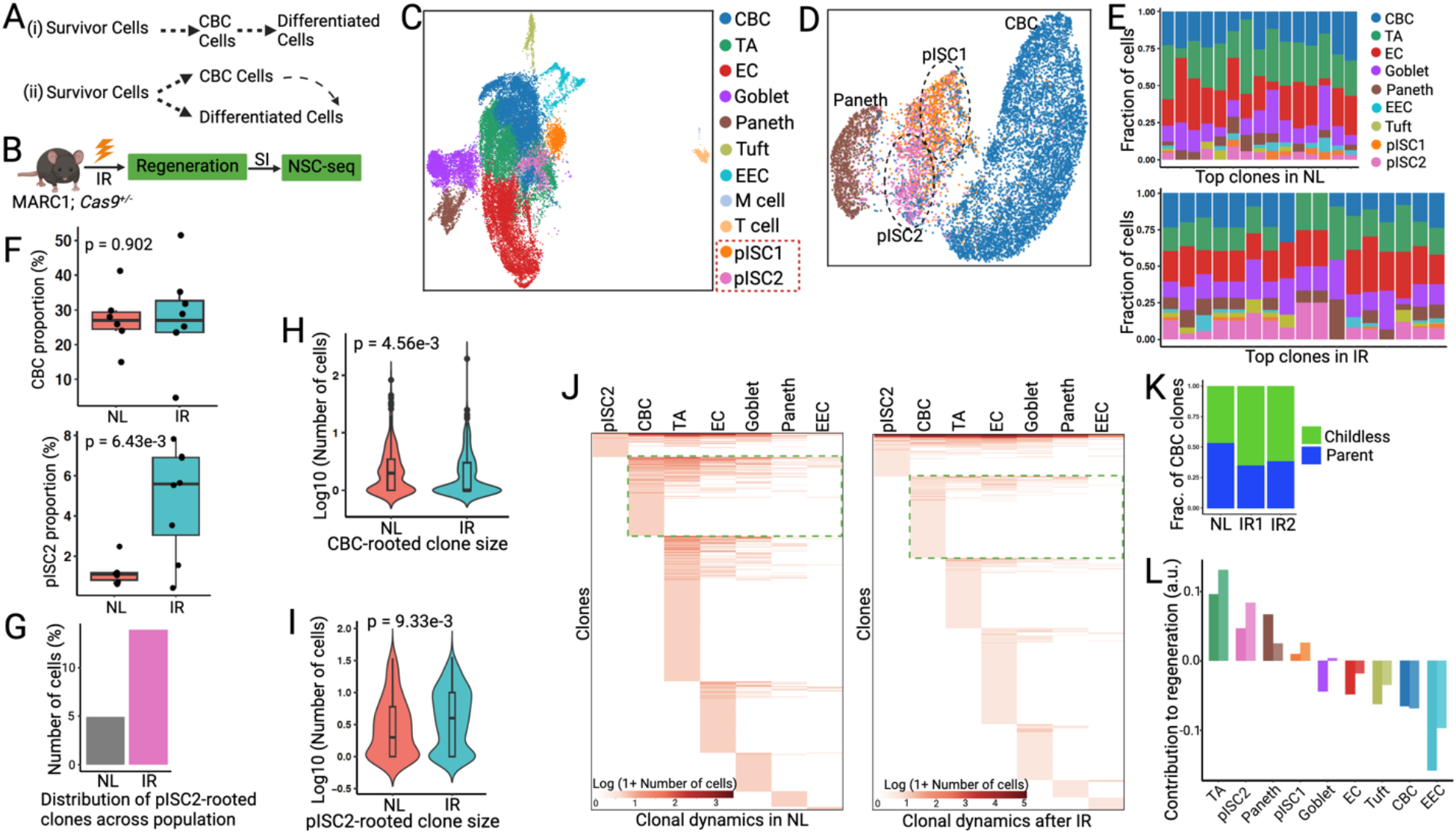
Clonal contributions of active and quiescent cell types in the irradiated intestinal epithelium. (A) Alternative models of cell-mediated regeneration post-irradiation (IR). (B) Experimental outline using native single-guide RNA capture and sequencing (NSC-seq) to query the contribution of various intestinal stem cell (ISC) populations after IR in an unbiased fashion. (C) Uniform manifold approximation and projection (UMAP) embedding of NSC-seq data showing major intestinal cell populations. Cells are colored by annotated cell types. (D) UMAP sub-clustering of selected epithelial cell types. (E) Clonal distribution across cell types in homeostasis and regeneration. (F) CBC and persister (p)ISC proportions among epithelial cells in normal (NL) and irradiated (IR) conditions. (G) Shared proportion of pISC2-rooted clones across epithelial populations. (H-I) Violin plots represent CBC-and pISC2-rooted clone size. Box plots inside the violin show the median value (thick line), box edges represent the first and third quartiles and whiskers extend to the minimum and maximum within 1.5× interquartile range (IQR). P value from unpaired two-tailed t-test. (J) Clone distribution across cell types during homeostasis and regeneration. Green box highlights CBC-derived clones. Heat map color represents the number of cells found comprising a clone within a given cell type. (K) Fraction of CBC-derived parent and childless clones in NL and IR conditions. (L) Relative contribution to regeneration by different cell types (n =2). Y-axis represents the change in parent clone fraction after IR compared to normal. Positive values indicate active contribution, negative values indicate no notable contribution. See also Figure S1.

Notably, while recently identified revival stem cells (revSCs) were only detectable during IR-induced regeneration [9], pISCs can be detected in both the small intestinal and colonic epithelium under homeostatic conditions (Figures S1Q-S1S), implicating their role as a primordial reserve cell that can give rise to other populations. To define the pISC’s role as a progenitor versus a dedifferentiating population, we first examined a mouse model where the secretory lineage differentiation is blocked (*Lrig1*^*CreERT2/+*^; *Atoh1*^*fl/fl*^) [29]. pISCs were present in this model, indicating that they are not secretory progenitors (SP) (Figure S1T). Furthermore, gene expression analysis did not support pISCs as label-retaining Paneth cells or enteroendocrine progenitors [30] (Figure S1U). We also assessed the expression of absorptive progenitor markers [31] and found no specific association with pISCs. Finally, we compared the pISC gene signature to that of the recently reported revSC population [9] and found no significant overlap between the two populations (Figure S1V). Together, these findings suggest that pISCs represent a population distinct from revSCs [9], labeling-retaining progenitors [30], or Fgfbp1^+^ upper crypt cells [15], which are found at homeostasis and are mobilized to contribute to other cell populations in response to damage.

Using an orthogonal bulk RNA-seq approach, we identified a quiescent subset of Lrig1+ cells [32] (Lrig1-Apple-mid) within mouse intestinal crypts. One of the anti-proliferative genes expressed in these cells was *Tob2* (Figures S1W-S1Y), which also emerged as part of the gene signature discovered in pISCs (Figure S1Z). TOB2 is a member of the TOB/BTG family of anti-proliferative (APRO) proteins that regulate global mRNA levels as part of CCR4-Not complex [33] and suppresses cellular proliferation [34]. Other members of this family, such as Tob1, are expressed by quiescent T cells [35] and osteoblasts [36], while other family members, Btg1 and Btg2, are known to be expressed by quiescent neural stem cells [37]. Indeed, overexpressing TOB2 in HEK293 cells significantly reduced cellular growth (Figure S1Z’), supporting its anti-proliferative role. Furthermore, we observed an inverse relationship between Tob2 expression and global mRNA expression across cell types (Figure S1E), likely reflecting the deadenylating function of the TOB/BTG family of proteins. Collectively, these findings suggest that Tob2 serves as a functional marker for the normally quiescent pISC population.

### Tob2 marks the pISC population

To validate Tob2 as a marker of the pISC population, we generated Tob2-CreERT2 mice in which endogenous Tob2 expression is unaffected (Figure 2A). One day after tamoxifen (TAM) induction, we observed infrequent lineage labeling, primarily labeling of single quiescent cells near the base of both small intestinal and colonic crypts (Figures 2B-2D, S2A, and S2B). Importantly, we confirmed that these labeled cells were not senescent and could move within the crypt over time (Figures S2C and S2D). Lineage tracing over 7 days revealed mostly small-sized clones; however, fully labeled crypt ribbons were observed, indicating Tob2^+^ cells possess stem potential and multiple states of activation during homeostasis (Figures 2E and S2E). Furthermore, Tob2^+^ cells gave rise to all differentiated cell types in both the small intestinal and colonic epithelium (Figure S2F). While overall lineage labeling and the presence of small clones diminished over time, the proportion of large clones increased, implying Tob2^+^ cells generate CBCs upon cell division in the unperturbed state (Figure 2F and S2G-S2I). Notably, the frequency of ribbons in the small intestine and colon derived from Tob2-CreERT2 mice do not show clonal extinction over time, in contrast to the 75% reduction of Lgr5-CreERT2-derived ribbons within 6 months [38], suggesting that single-labeled Tob2^+^ cells could expand into ribbons and compensate for the typical loss of ribbons due to neutral drift (Figure 2G) [39]. We also identified ribbons of labelled cells 7.5 months after a single tamoxifen induction (Figure S2J). These findings suggest that the Tob2+ cell population sustains long-term clonal output, a characteristic feature of long-term stem cells [40].

**Figure 2.**
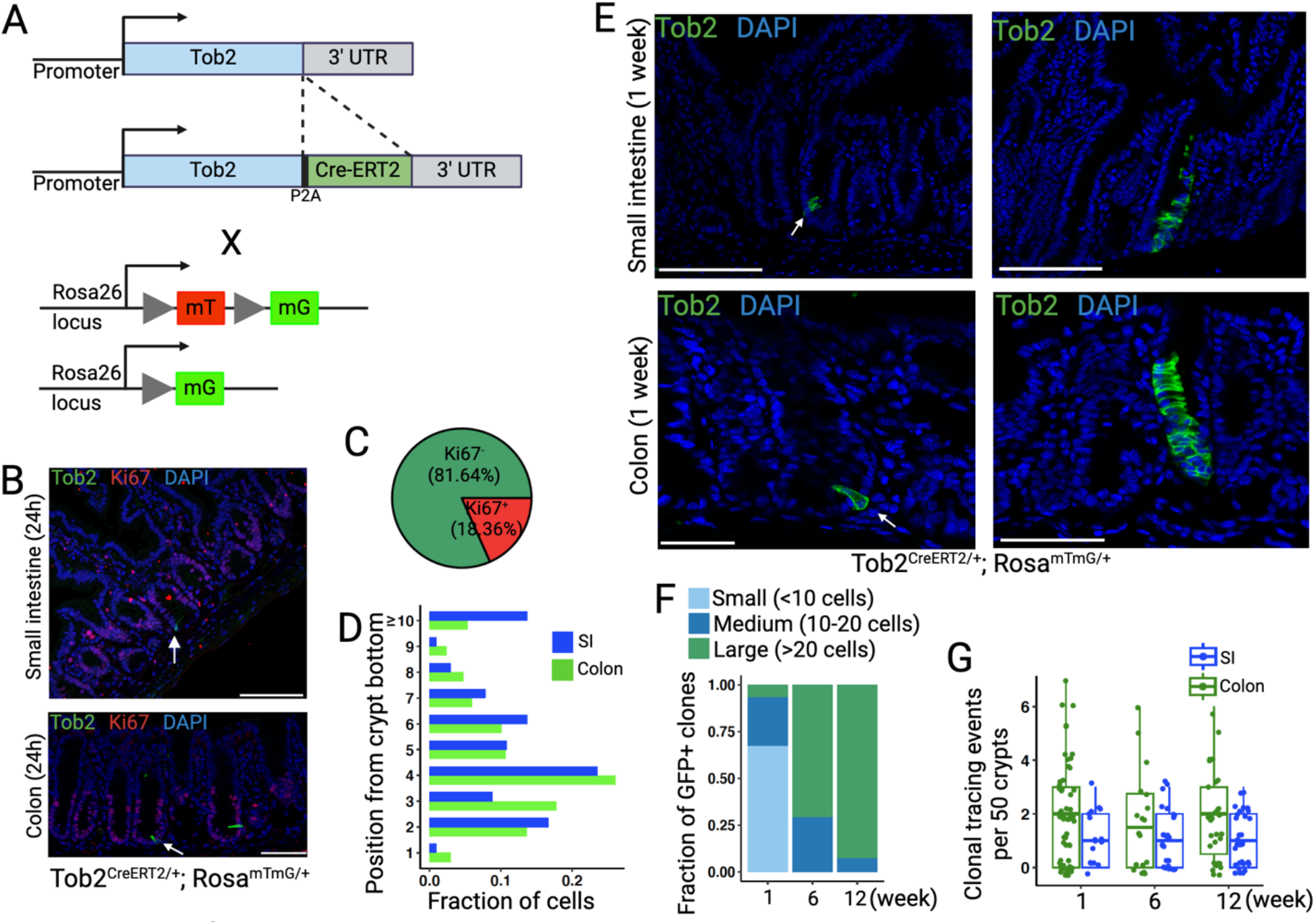
Tob2^+^ pISCs in the homeostatic intestinal epithelium. (A)Schematic for labelling Tob2^+^ cells and their progenies with GFP in a murine model. (B) Section of small intestine and colon 24h after TAM induction and imaged for GFP (green), Ki67 (red), and DAPI (blue) show examples of rare GFP^+^ cells that are Ki67^-^ (n = 3 independent experiments). (C) The proportion of GFP^+^ cells that are Ki67^+^ after 24h. (D) GFP^+^ clone position after 24h in small intestine (duodenum) and colon. (E) Sections of small intestine and colon 7 days after TAM induction (3 days pulse + 4 days chase) showing GFP^+^ clones. Representative images include small clones (left) and expanded ribbons (right) (n = 3 independent experiments). (F) GFP^+^ clone sizes quantified at 1-, 6-, and 12-week chase, plotted as fraction of total clones from three mice. (G) The frequency of ribbons quantified (n = 3) at 1-, 6-, and 12-week time points in small intestine (duodenum) and colon. Box plots show the median value, with edges representing the first and third quartiles and whiskers extend to the minimum and maximum within 1.5× IQR. See also Figure S2.

We then examined Tob2-CreERT2 lineage labeling across various tissues and found labeling in quiescent cell types, such as hepatocytes and skeletal muscle cells (Figure S2K). To better characterize these cells specifically in the GI tract, we flow sorted Tob2-labeled cells from both small intestine and colon, followed by scRNA-seq, which revealed three distinct clusters: pISC1, pISC2, and CBC-like cells (Figures S2L and S2M). Notably, pISC1 and pISC2 clusters co-embedded with previously identified pISCs, confirming that Tob2-CreERT2 marks the pISC population. The CBC-like cluster exhibited elevated biosynthetic capacity and was marked by Polr1a expression, a gene linked to ribosomal DNA transcription. Polr1a-expressing cells have been shown to maintain the stem cell hierarchy with Lgr5^+^ cells in a colorectal cancer model [41]. Although the CBC-like cell cluster shared some transcriptional similarities with Lgr5^+^ CBCs, we found them to be epigenetically distinct with higher chromatin accessibility than CBCs (Figure S3). Lastly, we found that sorted pISCs formed organoids in Matrigel culture, though at a lower efficiency than Lgr5+ cells (Figures S2N-S2P). Collectively, these findings suggest that Tob2 marks pISC populations, which exist in different states of activation during homeostasis in the intestinal epithelium.

### Origin of adult pISCs

Given the importance of chromatin states in revealing cell state heterogeneity and developmental potential, we simultaneously profiled gene expression and chromatin accessibility in sorted Tob2^+^ and Lgr5^+^ cells using the 10X multiome platform (Figures 3A-3D, S3A, and S3B). We found that these two populations were transcriptionally and epigenetically distinct. We identified differential chromatin accessibility between pISC1/2, CBC-like, and CBCs with a gradient among these populations in group 2 peaks (Figure 3E), which correlated with a Tob2 expression gradient (Figure 3C). Overall, Tob2^+^ cells displayed higher chromatin accessibility compared to Lgr5^+^ cells, particularly in enhancer regions (Figures 3F and S3C-S3E). CBC-like cells also showed greater accessibility than CBCs, arguing against the possibility that these cells are SPs, as SPs exhibit lower chromatin accessibility than CBCs (Figure S3C) [42]. No accessibility bias was observed in genes specific to secretory or absorptive progenitors (Figure S3F), supporting the conclusion that pISCs have not yet committed to a differentiated lineage. Pseudotime trajectory analysis [43] revealed a gradient between CBC-like and CBC populations, highlighted by unspliced unique Lgr5 transcript count residuals (Figure S3G). In CBC-like cells, Lgr5 transcripts retained a high ratio of introns (Figure S3H), indicating a transition from CBC-like (early) to CBC (late), analogous to the early-to-late transition seen in enteroendocrine differentiation [44]. We also identified distinct transcription factor motifs and regulatory profiles between Tob2^+^ and Lgr5^+^ cells (Figures S3I and S3J). Interestingly, Tob2^+^ cells exhibited higher expression of fetal gene signatures, as well as higher chromatin accessibility of reported fetal genes than Lgr5^+^ cells (Figures 3G and S3K-M) [45]. Using a curated set of fetal-specific intestinal peaks [46], we found increased accessibility in Tob2^+^ cells, following a similar Tob2 expression gradient as noted before (Figures 3H and S3N). Moreover, adult pISCs showed a higher mosaic fraction (MF) of early embryonic mutations (EEM), indicating their origin during early fetal development and persistence in adulthood (Figure S3O) [28]. Profiling fetal epithelial cells using NSC-seq revealed that Tob2^+^ cells were nearly twice as abundant as Lgr5^+^ cells at E14.5 in the gut epithelium, with little overlap between the two populations (Figures 3I, 3J, and S3P). Reported fetal genes such as Clu and Emp1 [45], were expressed in a similar fashion to Tob2 at E14.5, suggesting that Tob2 could also be a fetal marker gene (Figure S3Q). Moreover, fetal Tob2^+^ cells showed higher CytoTRACE scores and transcriptome counts compared to Tob2^-^ cells, characteristic of fetal stem cells (Figures S3R and S3S). Barcode analysis also confirmed their independent contribution to epithelial development at this early embryonic stage (Figure 3K). Lineage labeling of Tob2^+^ cells (*Tob2*^*CreERT2/+*^; *Rosa*^*mTmG/+*^) during embryogenesis further demonstrated their contribution to epithelial development at the E12.5 and E14.5 stages (Figures 3L-O and S3T-S3V). Unlike Lgr5-CreERT2 mice, which showed random labeling in the midgut at early embryonic stages [47], Tob2-CreERT2 mice demonstrated consistent and uniform lineage tracing at these early embryonic stages. Together, these data establishes that fetal Tob2^+^ cells act as intestinal stem cells contributing to early epithelial development. Given the limited contribution of fetal Lgr5^+^ cells to the adult ISC pool [2] and the mutual exclusivity of Tob2^+^ and Lgr5^+^ populations, our findings suggest that Tob2^+^ fetal cells are one of the populations that contributes to adult homeostatic ISCs.

**Figure 3.**
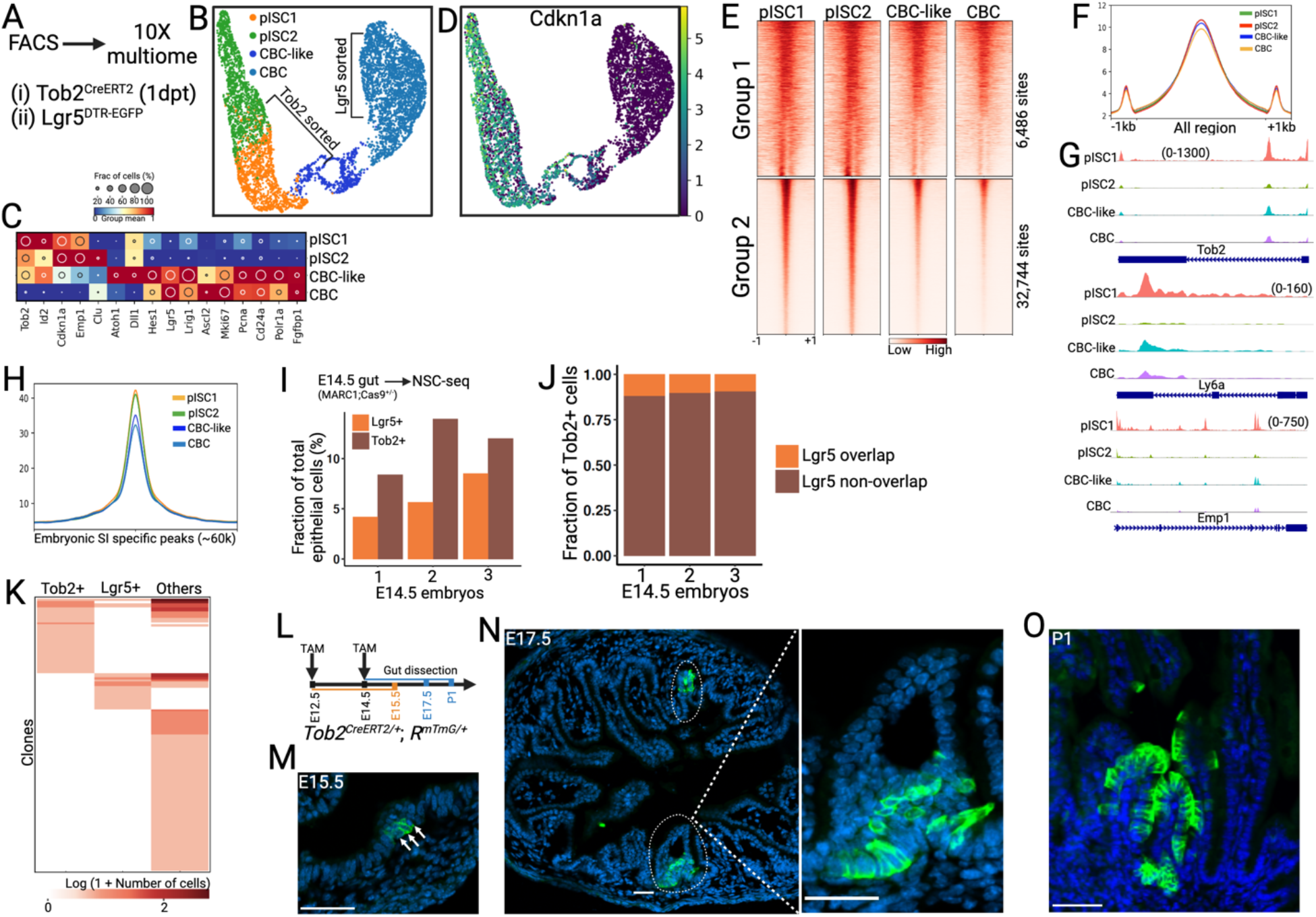
Origins of pISCs. (A) Schematic for simultaneous gene expression and chromatin accessibility experiments (10X multiome) of flow-sorted ISC populations. (B) UMAP embedding of transcriptomic data derived from sorted cell populations using multiome experiments. Cells are colored by annotated cell types. (C) Dot plots of cell type-specific marker genes. The size of the circle denotes the fraction of marker-positive cells, and color intensity indicates normalized group mean. (D) Normalized gene expression of Cdkn1a (p21). (E) ATAC-seq profile of top differentially expressed peaks across annotated cell types and K-means clustering. Group 2 peak regions lose accessibility in CBCs. (F) Average ATAC-seq signals across all peak regions in pISC1, pISC2, CBC-like and CBC. (G) Representative examples of ATAC-seq track plots of selected genes illustrate differential chromatin access. (H) Average ATAC-seq signals for E16.5 small intestine-specific peaks in adult pISC1, pISC2, CBC-like, and CBC. (I) Proportion of Tob2^+^ and Lgr5^+^ cells in the E14.5 intestinal epithelium. (J) Fraction of overlap between Tob2 and Lgr5 populations. (K) Distribution of clones across cell types at E14.5 intestinal epithelium. (L) Schematic for embryonic lineage tracing using Tob2-CreERT2 mouse line. (M) Section of embryonic gut tube after single dose of TAM induction at E12.5 and imaged for GFP (green) and DAPI (blue) at E15.5 showing examples of GFP^+^ clones. (N) Section of embryonic gut tube (small intestine) after single dose of TAM induction at E14.5 and imaged for GFP (green) and DAPI (blue) at E17.5 showing examples of GFP^+^ clones. Right, enlarged image of a crypt. (O) Section of small intestine shows fully labelled crypt-villus axis. Single dose of TAM induction at E14.5 and imaged for GFP (green) and DAPI (blue) at P1. Panel M-O imaged in at least 3 independent embryos. See also Figure S3.

### pISCs are essential to regeneration

To assess the regenerative capacity of adult pISCs, we lineage-labeled pISCs using Tob2-CreERT2 and induced epithelial damage with IR. Five days post-IR, we observed an increase in lineage-labeled cells per crypt (ribbons) in irradiated mice compared to non-irradiated controls, indicating the injury-responsive nature of pISCs (Figures S4A and S4B). To explore the role of Tob2 in regeneration, we generated mice with a germline targeted allele of *Tob2* (Figure S4C). Homozygous *Tob2* null mice (Tob2^-/-^) appeared normal at homeostasis; however, five days after IR (12 Gy), they exhibited severe epithelial destruction with edema and crypt loss in both the small intestine and colon (Figure 4A). All Tob2 null mice (n= 8) died by 10 days post-IR, indicating that *Tob2* is critical for survival after damage. In addition, we determined the role of Tob2^+^ pISCs in mouse models by genetic ablation of pISCs upon tamoxifen treatment (*Tob2*^*CreERT2/+*^; *Rosa*^*LSL- DTA/+*^). Five days post-tamoxifen, no notable epithelial morphological changes were detected (data not shown), suggesting that pISCs are dispensable under homeostatic conditions. However, after IR (12 Gy), pISC-ablated mice exhibited similar histological changes in the small intestine and colon as Tob2 null mice (Figures 4B and S4D), consistent with pISCs representing a major regenerative population in the crypts of both mouse small intestine and colon. We also performed scRNA-seq analysis on *Tob2* null (KO) and *Tob2* heterozygous (Het) mouse intestines, identifying distinct transcriptional profiles in *Tob2* KO pISCs with a notable increase in mRNA content (Figures S4E and S4F). *Tob2* KO pISCs showed elevated epithelial differentiation and TGFβ signaling, along with decreased expression of genes involved in antimicrobial responses and glutathione metabolism (Figures S4G-S4I). Similar analyses in the colon also revealed increased cell cycle signaling in *Tob2* KO pISCs (Figure S4J). Taken together, these results suggest that Tob2 is a functional marker of pISCs, and this population is critical for survival following IR.

**Figure 4.**
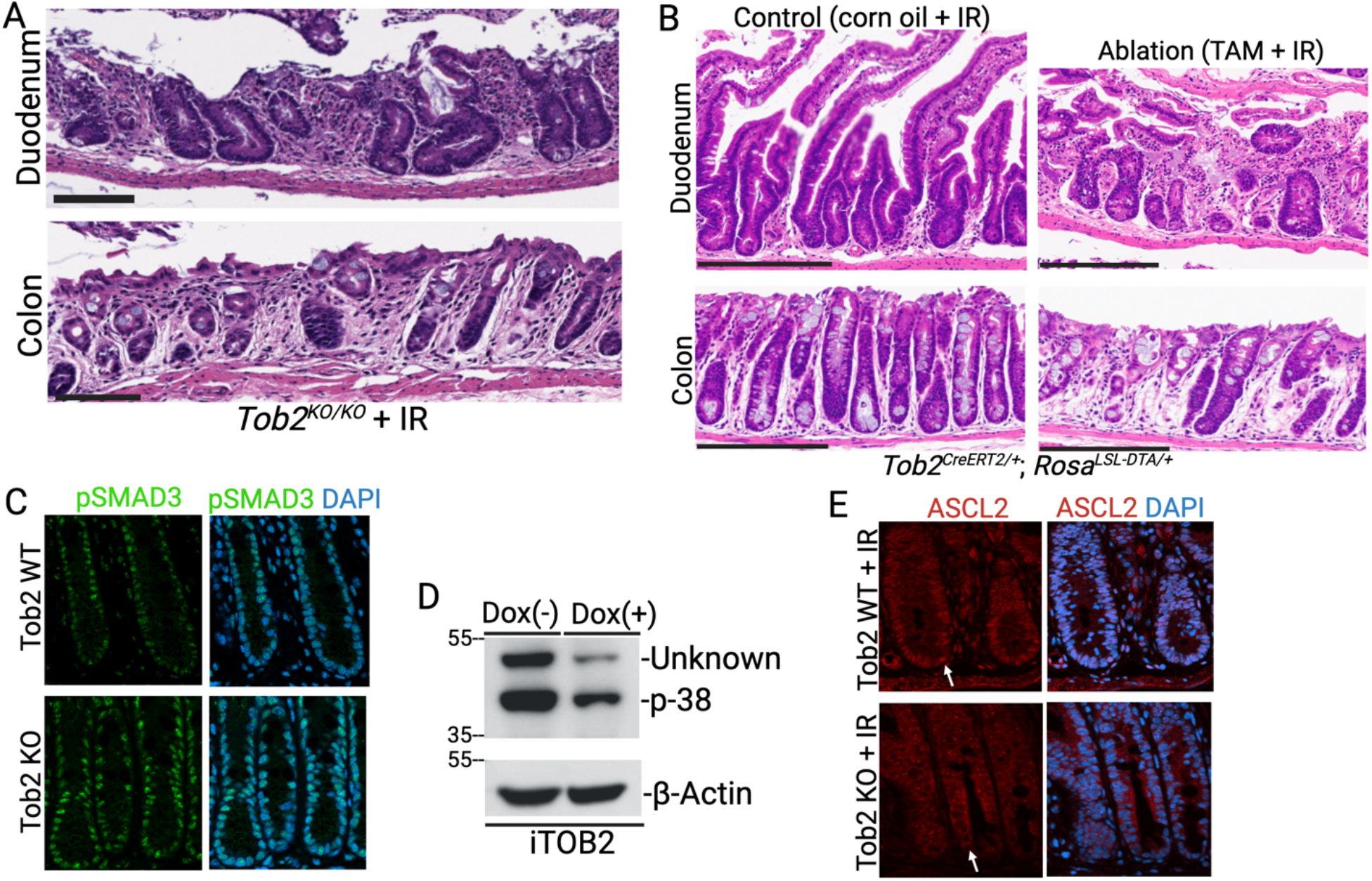
pISCs are essential for regeneration after IR. (A) Representative H&E section of duodenum and colon from Tob2^KO/KO^ mice after 5 days of IR. (B) Representative H&E staining of control (corn oil) and Tob2^+^ cells-ablated (TAM) duodenum and colon after 5 days of IR. Cre activation induces death of Tob2^+^ cells in this genetic model. Panel A-B are representative images from at least 3 independent mice. (C) Immunofluorescent staining of Phospo-SMAD3 (green) and DAPI (blue) in Tob2 WT and KO colonic crypts. (D) TOB2 overexpression in HEK293 cells reduces phosphorylated p-38 level by immunoblotting. Unknown represents unknown protein band. (E) Immunofluorescent staining of ASCL2 (red) and DAPI (blue) in Tob2 WT and KO colonic crypts after 5 days of IR. White arrows are indicating nuclear localization of ASCL2. See also Figure S4.

To dissect the mechanism of Tob2 in regulating stem cell properties *in vivo*, we generated Tob2 CRISPR KO MC-38 cell lines and implanted them into syngeneic C57BL/6 mice as xenografts. Tob2 KO led to increased invasive properties (Figures S4K-S4M), as well as increased TGFβ and Sonic Hedgehog (Shh) signaling (Figure S4N). Single-cell profiling of MC-38 xenografts also showed enhanced differentiation and cell cycle activity in Tob2 KO cells (Figures S4O and S4P). Furthermore, Tob2 KO cells exhibited elevated pSMAD2/3, but not pSMAD1/5, further implying that TOB2 regulates TGFβ signaling (Figure S4Q). This observation was also supported by increased pSMAD3 immunostaining in Tob2 KO crypts compared to Tob2 WT crypts (Figure 4C). Overexpression of TOB2 in HEK293 cells resulted in reduced phospho-p38 levels, supporting overall regulation of both canonical and non-canonical TGFβ signaling (Figures 4D and S4R). To further examine the function of TOB2, we performed yeast two-hybrid (Y2H) screens and found several interacting proteins, including ASCL2 (Table S1); the interaction between TOB2 and ASCL2 was confirmed in a yeast one-hybrid assay (Figure S4S). ASCL2 expression was increased during IR-induced regeneration (Figure S4T). We performed immunofluorescent staining of ASCL2 in TOB2 wild-type (WT) and KO mouse colonic tissues, which revealed impaired nuclear localization of ASCL2 in TOB2 KO colonic epithelium after IR (Figure 4E). Moreover, using publicly available datasets [48], we found increased expression of TGFβ target genes in the ASCL2 KO mouse intestine, similar to increased TGFβ signaling in the Tob2 KO condition, supporting a reciprocal regulation of TOB2 and ASCL2 (Figures S4U-W). Collectively, these findings suggest that TOB2 is crucial for maintaining the balance between TGFβ-mediated fetal differentiation [49] and ASCL2-mediated stemness [48], promoting a rapid regenerative response after injury.

### Human LGR5+ cells are quiescent

While the clonal behavior of human intestinal epithelial cells has been studied using organoids or xenograft models, their dynamics *in situ* remain unclear [18, 19]. Here, we investigated the lineage hierarchy and clonal behavior of the human colonic epithelium under homeostatic conditions by leveraging naturally occurring mitochondrial DNA mutations as lineage barcodes, combined with cell-state information for phylogenetic reconstruction. As part of the Human Tumor Atlas Network (HTAN) initiative, we generated scRNA-seq data from 65 normal human colonic biopsies obtained at the time of screening colonoscopy (Fig. 5A and Table S2). Lineage-informative mitochondrial variants (mtVars) were identified at the single-cell level using a previously validated approach (Figures S5A-S5F) [28]. Cells were annotated using gene expression profiles from both the proximal and distal colonic epithelium (Figures S5G and S5H). Overall cell types and their proportions aligned with previous reports [50], though the proportion of infiltrating lymphoid cells varied with age (Figures S5I and S5J). Within the STEM cluster, we identified a unique LGR5^+^PCNA/KI67^-^ subcluster (hereafter referred to as LGR5^+^ cells) that expressed BTG2, a member of the TOB/BTG family of APRO proteins, and the cell-cycle inhibitor p27 (CDKN1B) (Figures 5B, 5C, and S5K-S5M). While these cells exhibited a low proportion of G2M-phase activity, they surprisingly showed a high proportion of S-phase activity - a putative hallmark of reserve ISCs with replicative quiescence (Figure S5N) [51]. Additionally, fluorescence *in situ* hybridization (FISH) staining confirmed mutual exclusivity of KI67 and LGR5 expression (Figure 5D). Finally, this subcluster exhibited a similar mtVars burden to STEM-assigned cells (STEM cluster without a LGR5^+^ subcluster, as designated by STEM-associated gene expression), despite demonstrating a relatively high stemness potential (Figures 5E and S5O). These findings suggest that homeostatic LGR5^+^ cells in the human colon are predominantly quiescent in marked contrast to their murine Lgr5^+^ counterparts.

**Figure 5.**
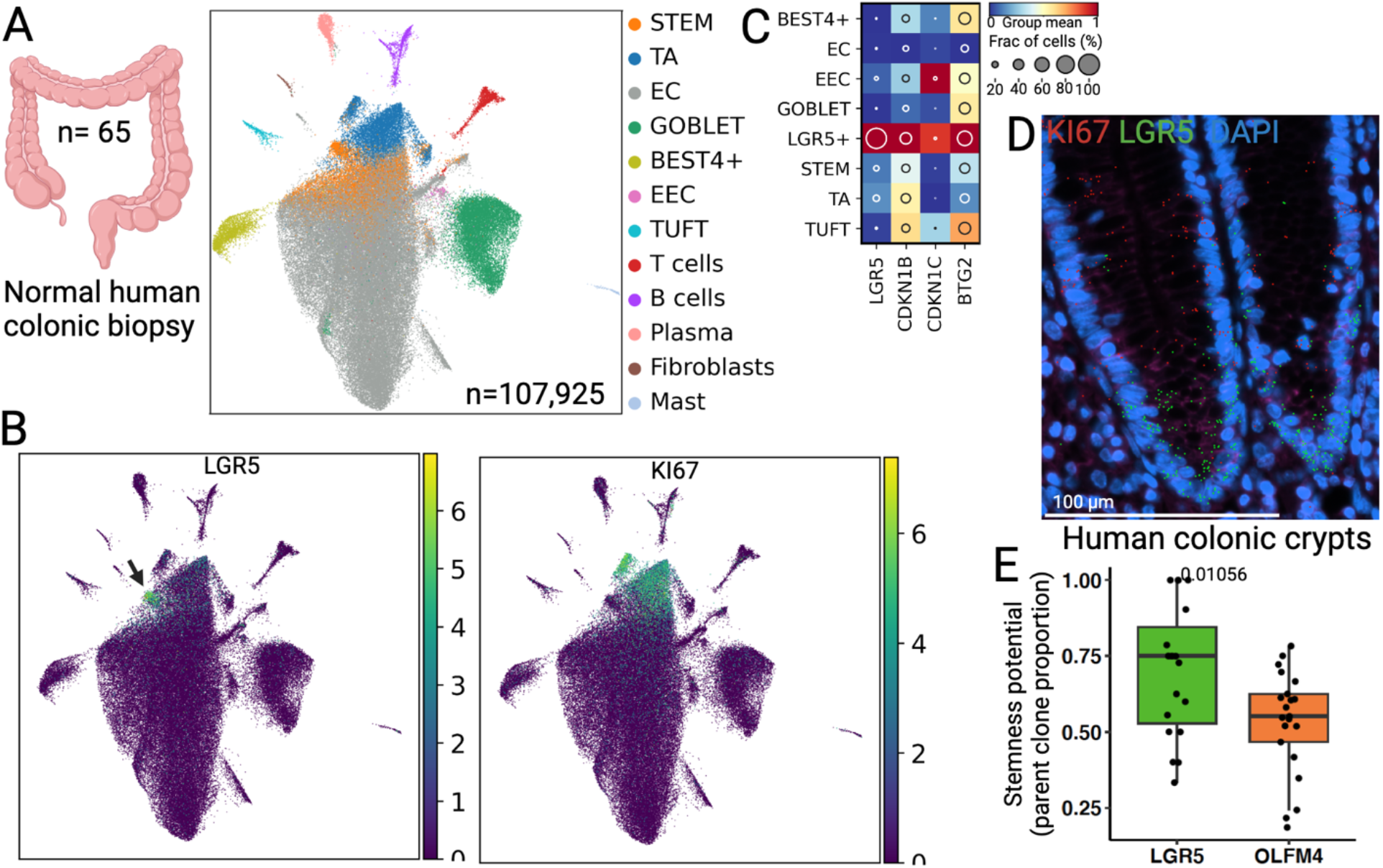
Single-cell analysis of human colonic epithelium. (A) Overview of scRNA-seq experiment from proximal (ascending) and distal (descending) colonic regions (left). UMAP embedding shows cell populations from scRNA-seq experiments (right). Cells are colored by annotated cell types. (B) Normalized gene expression of LGR5 (black arrow) and KI67. (C) Dot plots of selected marker genes across epithelial cell types. The size of the circle denotes the fraction of marker-positive cells, and color intensity indicates normalized group mean. High BTG2 and CDKN1B (p27) expression in LGR5^+^ cells. (D) FISH staining of LGR5 (green) and KI67 (red) in human colonic crypts. (E) Measurement of stemness potential across different cell types. See also Figure S5.

### Clonal architecture of the human colonic epithelium

Next, we observed that stem cells exhibited a significantly lower mtVars burden compared to differentiated colonocytes/enterocytes (hereafter referred to as ECs), with transit-amplifying (TA) cells displaying an intermediate mtVars burden (Figure 6A). This pattern suggests a clear lineage hierarchy, transitioning from stem cells to TA cells and subsequently to differentiated cells, analogous to the lineage hierarchy observed in the mouse intestine [1] and human hematopoietic system [24]. Additionally, analyses of stemness potential and rooted-clone size revealed significant differences between STEM/TA cells and ECs, further corroborating the existence of a lineage hierarchy in the human colonic epithelium that is consistent with murine models (Figures 6B and S6A). We then investigated regional differences in clonal behavior by comparing paired proximal and distal colonic tissues. Across samples, the distal colon displayed a higher mtVars burden than the proximal colon, indicative of a more rapid cellular turnover (Figure S6B and S6C). Of interest, despite of small sample size, individuals of African ancestry exhibited a higher rate of cellular turnover compared to Caucasian ancestry, particularly in the proximal colon over time (Figure S6D). This observation aligns with reports of enhanced replicative aging signatures among individuals of African ancestry, potentially reflecting racial disparities as previously documented [52, 53].

**Figure 6.**
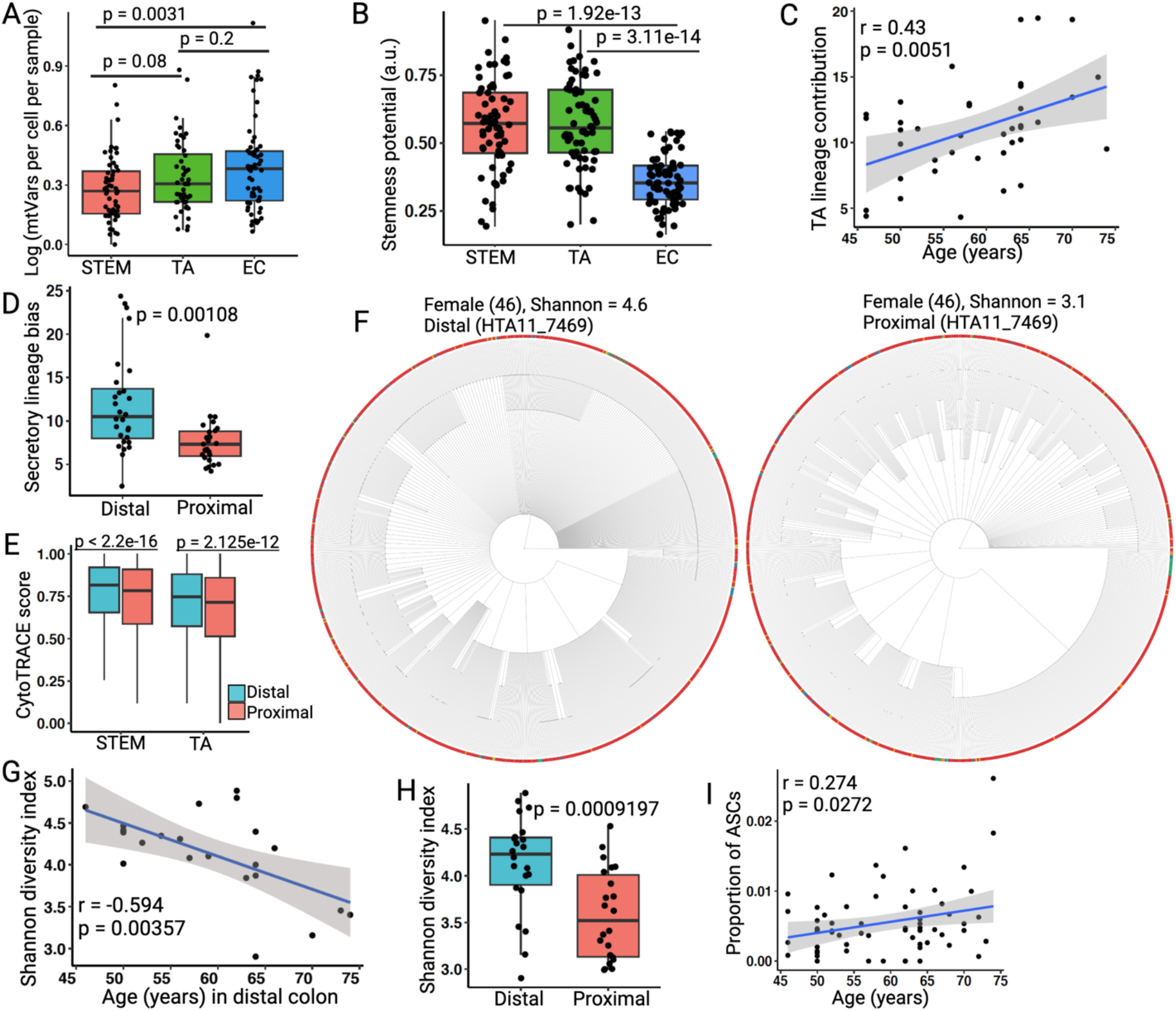
Clonal analysis of the homeostatic human colonic epithelium. (A) Measurement of mtVar burdens across selected cell types. (B) Measurement of stemness potential across different cell types. Stem potential was defined by the proportion of parent clones. See Supplemental Methods. (C) Pearson correlation analysis between TA lineage contribution and age. (D) Secretory lineage preference between proximal and distal colonic regions. (E) CytoTRACE scores between proximal and distal colonic tissues for selected cell types. (F) Reconstructed single-cell lineage tree from proximal and distal colonic regions. Leaf cells are colored by annotated cell-type colors, except EC is marked as red. Nodes are colored by dark gray. Each branch represents an independent mtVar event. Non-binary single-cell trees for other samples can be found in the NSC-seq GitHub page. (G) Pearson correlation analysis between Shannon diversity index of all cells and age in distal colon. (H) Measurement of Shannon diversity index between paired proximal and distal biopsies for all cells. (I) Pearson correlation analysis between proportion of adenoma-specific cells (ASCs) in normal biopsy and age. A,B,D,E,H, Box plots showed the median value (thick line), box edges represented the first and third quartiles and whiskers extend to the minimum and maximum within 1.5× IQR. P values were derived from unpaired two-tailed t-test. C,G,I, Correlations and P values (by F-test) and 95% confidence intervals (shaded area) are indicated. See also Figure S6.

Finally, we performed a clonal distribution analysis that revealed a significant increase in TA lineage preference over time, supporting an increasing contribution from TA cells, while stem cell dynamics remained relatively stable with age (Figures 6C, S6E, and S6F). Stem cells in the distal colon produced more secretory cells than the proximal colon regardless of age, indicating a secretory lineage bias in the distal colon across samples (Figure 6D) [50, 54]. Clonal fixation, estimated by stem cell replacement frequency as a proxy for increasing connectivity or clonal grouping within the stem cell population [24, 27], showed that the fraction of connected stem cells increased over time, supporting the fixation and expansion of clones across the epithelium (Figure S6G). However, the rate of connectivity increase was slow (∼0.8% per year), partially explaining why neutral drift in human colonic epithelium is much slower (>5 years) compared to the murine intestinal epithelium (∼2 months), despite a similar 3-7 days turnover [39, 55]. Since TA cells are short-lived, we did not observe increasing connectivity within TA cells over time (Figure S6H). Stem cell connectivity, mtVars burden, stemness potential, and CytoTRACE scores were higher in the distal compared to proximal colon (Figures 6E, S6I, and S6J), consistent with the higher cellular turnover in the distal colon. This observation may have implications for the greater number of premalignant adenomas as well as chromosomal instability (CIN) adenocarcinomas observed in the distal colon compared to proximal colon [56, 57]. Overall, we noticed clonal contributions beyond stem cells into differentiated cells, regardless of variability in age, gender, and race, and the existence of a lineage hierarchy in the human colonic epithelium *in situ* [19]. Single-cell lineage reconstruction showed distinct characteristics between the proximal and distal colonic epithelium with the distal colon showing larger clone sizes (Figures 6F and S6K). However, barcode diversity decreased over time in the distal colon but not in the proximal colon, suggesting bigger crypt patch size and a higher chance of spreading oncogenic mutations across the distal colonic epithelium, further emphasizing regional differences (Figures 6G, 6H, and S6L-N). Despite these distinctions, we found adenoma-specific cells (ASCs) [58] within most of the normal colonic tissue, albeit rare (∼0.05%), with a significant association with increasing age (Figure 6I). This observation aligns with findings of age-dependent abnormal cells in the human breast epithelium [59]. Collectively, these results reshape our understanding of clonal dynamics in the normal human colonic epithelium and highlight how aging influences clonal architecture and possibly cancer risk.

## DISCUSSION

Our findings uncover distinct cellular behaviors across species, such as the quiescent nature of human LGR5^+^ cells *in situ* in marked contrast to the rapid cycling of murine Lgr5^+^ cells. These observations underscore species-specific differences, highlighting the importance of studying native tissue dynamics and the need for human-relevant model systems. We speculate that the quiescent nature of human LGR5^+^ cells might represent an evolutionary adaptation to mitigate cancer risk in high-turnover tissues and to extend lifespan, which is inconsistent with Peto’s paradox [60]. This quiescence likely slows the fixation and spread of oncogenic mutations, despite the presence of rare adenoma-specific cells in the normal epithelium [27, 55]. Future studies comparing clonal fixation under paired homeostatic and oncogenic mutational contexts will expand our understanding of human cancer development. Our data reveal regional differences in clonal dynamics between the proximal and distal human colonic epithelium, which may have implications for tissue physiology and disease prevalence, including cancer risk [61]. From this work, aging emerged as a critical factor influencing clonal diversity in the human colon. However, our study lacked longitudinal sampling from the same individuals, which could further elucidate how clonal architecture evolves over time in the human colonic epithelium.

The NSC-seq approach enabled us to assess the clonal potency and gene expression of various cell types simultaneously, an approach that overcomes the limitations of traditional marker-based lineage mapping in murine models [28]. Consistent with previous studies, we confirmed the contribution of Paneth cells to intestinal regeneration [8]. However, different types of damage may activate distinct regenerative populations. It was surprising to find the dramatic impact of genetically ablating the rare pISC population, which, although dispensable during homeostasis, severely impaired the overall regenerative capacity of the intestinal epithelium post-IR. This suggests that dysregulation of TGFβ signaling and impaired nuclear localization of ASCL2 may hinder CBC replenishment during regeneration, compromising intestinal plasticity and stemness, therefore supporting an indispensable role of pISCs and CBCs in regeneration [6]. These data suggest a model where intestinal regeneration relies on a collective hierarchical response rather than a single resident cell type required for regeneration.

We also provide evidence that pISCs specified during early embryogenesis contribute to gut development and persist into adulthood and after IR. Our clonal potency analysis supports a limited contribution from embryonic Lgr5^+^ precursors in gut development, similar to previous reports [2]. Interestingly, embryonic Lgr5^+^ cells play a region-specific role, with a smaller contribution to midgut development in comparison to the more pronounced role in hindgut development [47]. Although the contribution of fetal Lgr5^+^ precursors to the adult ISC pool is limited, our findings reveal fetal Tob2^+^ pISCs as an additional contributing population, highlighting the molecular and functional heterogeneity of the early embryonic intestinal epithelium. These data support a dynamic quiescent-to-active continuum in the adult mouse intestinal epithelium, where pISCs replenish CBCs (Figure 7A). Additionally, the recent observation that CBCs can be generated from Fgfbp1+ upper crypt zone represents a notable advancement [15]; however, we found Fgfbp1 is expressed in the TA/upper crypt zone and CBC-like cells but not in pISC1/2 cell population. This dynamic underscores the idea that pISCs serve as a “savings account” to replenish the actively dividing “checking account” nature of CBCs after exhaustion and their possible co-existence in a highly proliferative niche [11]. It is tempting to speculate that Tob2^+^p21^+^ pISCs in mice are a likely orthologue to LGR5^+^BTG2^+^p27^+^ populations in humans, given their similar quiescent state and role in regeneration (Figure 7B) [18].

**Figure 7.**
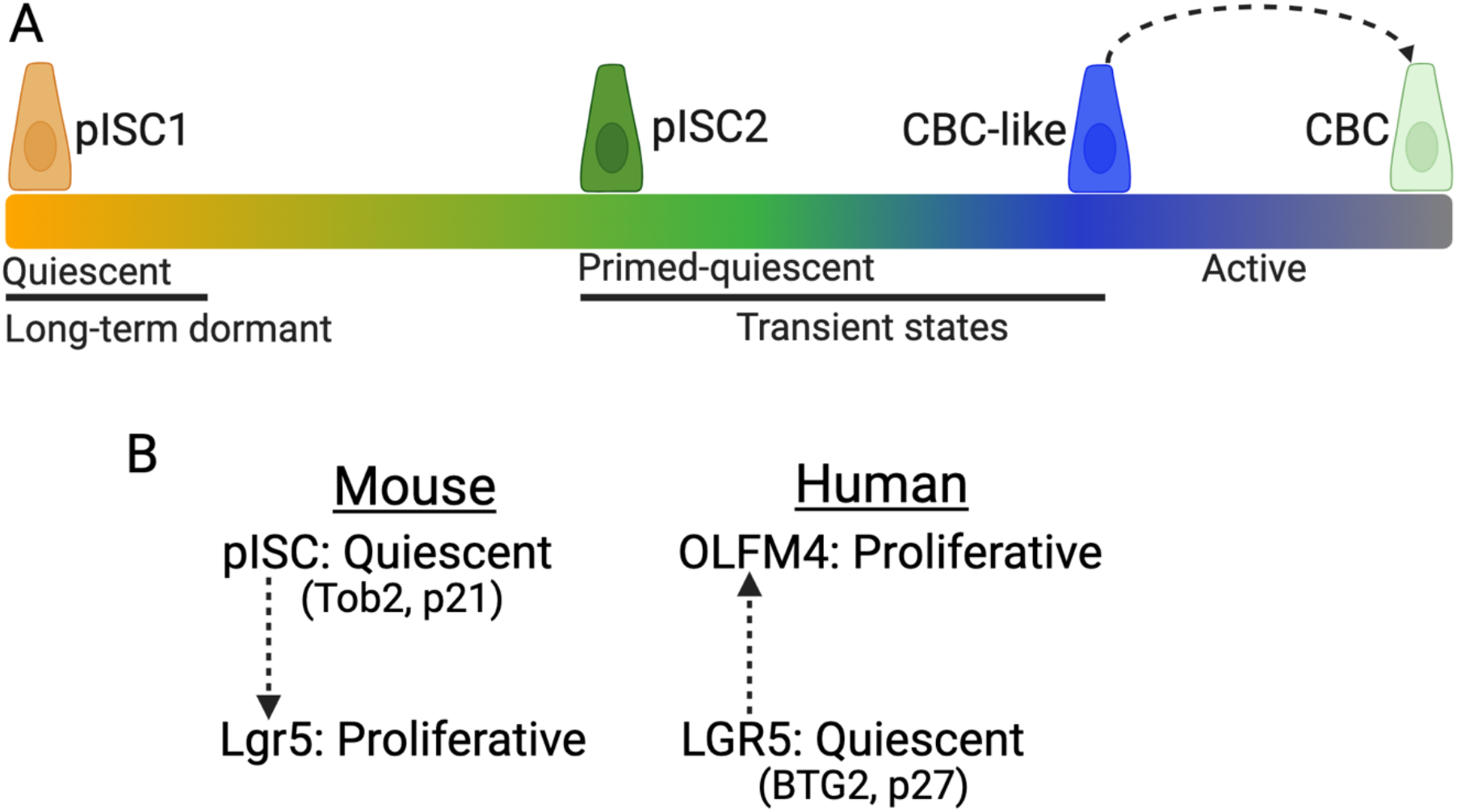
Proposed model for the mammalian intestinal stem cell regulation. (A) Schematic representation of mouse pISC populations representing quiescent-to-active continuum. (A) Summary of quiescent and active states between mouse and human intestinal crypts. The quiescent state is achieved in mammalian intestinal crypts by homologous sets of proteins, Tob2 and p21 in pISCs in mouse and BTG2 and p27 in LGR5^+^ cells in human.

Beyond intestinal biology, these findings highlight a broader paradigm of tissue plasticity and repair across organs. Unbiased lineage tracing approaches provide a framework for studying similar processes in other high-turnover tissues, potentially guiding therapeutic strategies to enhance regeneration in disease and injury contexts. While existing methods for lineage analysis with human mitochondrial mutations require specialized enrichment [24, 27], our approach directly utilizes scRNA-seq datasets, offering a scalable framework for clonal potency analysis and phylogenetic reconstruction. Despite the limited resolution without targeted mitochondrial read enrichment, this framework broadens the applicability of spontaneous mitochondrial mutations for dissecting cellular dynamics of human tissues in health and disease.

## Acknowledgements

This publication is part of the HTAN (Human Tumor Atlas Network) consortium paper package. The authors wish to thank the study participants and funding support by HTAN U2CCA233291 (to R.J.C., K.S.L., and M.J.S.), TBEL U54CA274367 (to R.J.C., K.S.L., and M.J.S.), R35CA197570 and P50CA236733 (to R.J.C.), R01DK103831 (to K.S.L.), K07CA122451 (to M.J.S.). R.J.C. acknowledge the generous support of the Nicholas Tierney Memorial GI Cancer Fund. We thank current and former members of the Lau and Coffey laboratories (especially Jumpei Kondo, Won Jae Huh, and Hiroaki Niitsu) for assistances with animal housing, tissue imaging, and single-cell data collection. Cores used by this study included Survey and Biospecimen Shared Resource, TPSR (U24DK059637), VANTAGE, REDCap (UL1TR000445), and Vanderbilt Genome Editing Resource (VGER). 1cellbio and RAN biotechnologies helped to synthesis the custom hydrogel beads. We use BioRender for drawing many schematics in this study. We apologize in advance to those we have failed to acknowledge due to space constraints.

## Authors’ contributions

Conceptualization was the responsibility of M.I. Supervision was undertaken by M.I., K.S.L. and R.J.C. Data curation was carried out by M.I., M.E.B., Y.Y., A.J.S., Y.X., J.N.H, P.Z., Z.C., N.M., F.R., M.A.R.-S., K.S.L. and R.J.C. Formal analysis was the responsibility of M.I., Y.Y., and M.B. Methodology was the responsibility of M.I., M.E.B., and Y.Y. Project administration was carried out by M.I., M.E.B., A.J.S., K.S.L and R.J.C. Resources were the responsibility of M.I., Q.L., M.J.S., R.J.C. and K.S.L. Software was the responsibility of M.I., Y.Y., and K.S.L. Validation was carried out by M.I., M.E.B., Y.Y., K.S.L., R.J.C. Visualization was undertaken by M.I., M.E.B., and Y.Y. Original draft was written by M.I. Draft edited by M.I., S.E.G., K.S.L., and R.J.C. Reviewing of the text was performed by M.I., M.E.B., Y.Y., A.J.S., Y.X., J.N.H., P.Z., Z.C.,N.T., S.E.G., N.O.M., F.R., M.A.R.-S., Q.L., J.L.F, K.S.L., and R.J.C.

## Data and code accessibility

Human data have been deposited to the HTAN Data Coordinating Center Data Portal at the National Cancer Institute: https://data.humantumoratlas.org/ (under the HTAN Vanderbilt Atlas). Mouse data deposited in GEO: GSE285517. Reviewer token: *******. NSC-seq data analysis pipeline reported in GitHub: https://github.com/Ken-Lau-Lab/NSC-seq. Additional reconstructed single-cell lineage trees for both human and mouse intestine will be made available on our GitHub repository. Newly generated mouse lines will be available upon request to Robert Coffey.

## References

1. Barker, N., et al., Identification of stem cells in small intestine and colon by marker gene Lgr5. Nature, 2007. 449(7165): p. 1003–7.

2. Guiu, J., et al., Tracing the origin of adult intestinal stem cells. Nature, 2019. 570(7759): p. 107–111.

3. Tao, S., et al., Wnt activity and basal niche position sensitize intestinal stem and progenitor cells to DNA damage. Embo j, 2015. 34(5): p. 624–40.

4. Potten, C.S., Extreme sensitivity of some intestinal crypt cells to X and gamma irradiation. Nature, 1977. 269(5628): p. 518–21.

5. Barker, N., Adult intestinal stem cells: critical drivers of epithelial homeostasis and regeneration. Nat Rev Mol Cell Biol, 2014. 15(1): p. 19–33.

6. Metcalfe, C., et al., Lgr5+ stem cells are indispensable for radiation-induced intestinal regeneration. Cell Stem Cell, 2014. 14(2): p. 149–59.

7. Tian, H., et al., A reserve stem cell population in small intestine renders Lgr5-positive cells dispensable. Nature, 2011. 478(7368): p. 255–9.

8. Yu, S., et al., Paneth Cell Multipotency Induced by Notch Activation following Injury. Cell Stem Cell, 2018. 23(1): p. 46-59.e5.

9. Ayyaz, A., et al., Single-cell transcriptomes of the regenerating intestine reveal a revival stem cell. Nature, 2019. 569(7754): p. 121–125.

10. Yan, K.S., et al., Intestinal Enteroendocrine Lineage Cells Possess Homeostatic and Injury-Inducible Stem Cell Activity. Cell Stem Cell, 2017. 21(1): p. 78-90.e6.

11. Li, L. and H. Clevers, Coexistence of quiescent and active adult stem cells in mammals. Science, 2010. 327(5965): p. 542–5.

12. Kretzschmar, K. and F.M. Watt, Lineage tracing. Cell, 2012. 148(1-2): p. 33–45.

13. Woodworth, M.B., K.M. Girskis, and C.A. Walsh, Building a lineage from single cells: genetic techniques for cell lineage tracking. Nat Rev Genet, 2017. 18(4): p. 230–244.

14. Muñoz, J., et al., The Lgr5 intestinal stem cell signature: robust expression of proposed quiescent ‘+4’ cell markers. Embo j, 2012. 31(14): p. 3079–91.

15. Capdevila, C., et al., Time-resolved fate mapping identifies the intestinal upper crypt zone as an origin of Lgr5+ crypt base columnar cells. Cell, 2024. 187(12): p. 3039-3055.e14.

16. Yan, K.S., et al., The intestinal stem cell markers Bmi1 and Lgr5 identify two functionally distinct populations. Proc Natl Acad Sci U S A, 2012. 109(2): p. 466–71.

17. Ramadan, R., et al., Intestinal stem cell dynamics in homeostasis and cancer. Trends Cancer, 2022. 8(5): p. 416–425.

18. Ishikawa, K., et al., Identification of Quiescent LGR5(+) Stem Cells in the Human Colon. Gastroenterology, 2022. 163(5): p. 1391-1406.e24.

19. Sugimoto, S., et al., Reconstruction of the Human Colon Epithelium In Vivo. Cell Stem Cell, 2018. 22(2): p. 171-176.e5.

20. Kalhor, R., et al., Developmental barcoding of whole mouse via homing CRISPR. Science, 2018. 361(6405).

21. Chan, M.M., et al., Molecular recording of mammalian embryogenesis. Nature, 2019. 570(7759): p. 77–82.

22. Bowling, S., et al., An Engineered CRISPR-Cas9 Mouse Line for Simultaneous Readout of Lineage Histories and Gene Expression Profiles in Single Cells. Cell, 2020. 181(6): p. 1410-1422.e27.

23. Ludwig, L.S., et al., Lineage Tracing in Humans Enabled by Mitochondrial Mutations and Single-Cell Genomics. Cell, 2019. 176(6): p. 1325-1339.e22.

24. Weng, C., et al., Deciphering cell states and genealogies of human haematopoiesis. Nature, 2024. 627(8003): p. 389–398.

25. Coorens, T.H.H., et al., Extensive phylogenies of human development inferred from somatic mutations. Nature, 2021. 597(7876): p. 387–392.

26. Baker, A.M., et al., Quantification of crypt and stem cell evolution in the normal and neoplastic human colon. Cell Rep, 2014. 8(4): p. 940–7.

27. Gutierrez-Gonzalez, L., et al., Analysis of the clonal architecture of the human small intestinal epithelium establishes a common stem cell for all lineages and reveals a mechanism for the fixation and spread of mutations. J Pathol, 2009. 217(4): p. 489–96.

28. Islam, M., et al., Temporal recording of mammalian development and precancer. Nature, 2024. 634(8036): p. 1187–1195.

29. Banerjee, A., et al., Succinate Produced by Intestinal Microbes Promotes Specification of Tuft Cells to Suppress Ileal Inflammation. Gastroenterology, 2020. 159(6): p. 2101-2115.e5.

30. Buczacki, S.J., et al., Intestinal label-retaining cells are secretory precursors expressing Lgr5. Nature, 2013. 495(7439): p. 65–9.

31. Tetteh, P.W., et al., Replacement of Lost Lgr5-Positive Stem Cells through Plasticity of Their Enterocyte-Lineage Daughters. Cell Stem Cell, 2016. 18(2): p. 203–13.

32. Powell, A.E., et al., The pan-ErbB negative regulator Lrig1 is an intestinal stem cell marker that functions as a tumor suppressor. Cell, 2012. 149(1): p. 146–58.

33. Chen, C.A., et al., Tob2 phosphorylation regulates global mRNA turnover to reshape transcriptome and impact cell proliferation. Rna, 2020. 26(9): p. 1143–1159.

34. Winkler, G.S., The mammalian anti-proliferative BTG/Tob protein family. J Cell Physiol, 2010. 222(1): p. 66–72.

35. Tzachanis, D., et al., Tob is a negative regulator of activation that is expressed in anergic and quiescent T cells. Nat Immunol, 2001. 2(12): p. 1174–82.

36. Yoshida, Y., et al., Negative regulation of BMP/Smad signaling by Tob in osteoblasts. Cell, 2000. 103(7): p. 1085–97.

37. Adusumilli, V.S., et al., ROS Dynamics Delineate Functional States of Hippocampal Neural Stem Cells and Link to Their Activity-Dependent Exit from Quiescence. Cell Stem Cell, 2021. 28(2): p. 300-314.e6.

38. Li, N., et al., Single-cell analysis of proxy reporter allele-marked epithelial cells establishes intestinal stem cell hierarchy. Stem Cell Reports, 2014. 3(5): p. 876–91.

39. Lopez-Garcia, C., et al., Intestinal stem cell replacement follows a pattern of neutral drift. Science, 2010. 330(6005): p. 822–5.

40. Cheng, H. and C.P. Leblond, Origin, differentiation and renewal of the four main epithelial cell types in the mouse small intestine. V. Unitarian Theory of the origin of the four epithelial cell types. Am J Anat, 1974. 141(4): p. 537–61.

41. Morral, C., et al., Zonation of Ribosomal DNA Transcription Defines a Stem Cell Hierarchy in Colorectal Cancer. Cell Stem Cell, 2020. 26(6): p. 845-861.e12.

42. Jadhav, U., et al., Dynamic Reorganization of Chromatin Accessibility Signatures during Dedifferentiation of Secretory Precursors into Lgr5+ Intestinal Stem Cells. Cell Stem Cell, 2017. 21(1): p. 65-77.e5.

43. La Manno, G., et al., RNA velocity of single cells. Nature, 2018. 560(7719): p. 494–498.

44. Gehart, H., et al., Identification of Enteroendocrine Regulators by Real-Time Single-Cell Differentiation Mapping. Cell, 2019. 176(5): p. 1158-1173.e16.

45. Mustata, R.C., et al., Identification of Lgr5-independent spheroid-generating progenitors of the mouse fetal intestinal epithelium. Cell Rep, 2013. 5(2): p. 421–32.

46. Francis, R., et al., Gastrointestinal transcription factors drive lineage-specific developmental programs in organ specification and cancer. Sci Adv, 2019. 5(12): p. eaax8898.

47. Zhao, L., W. Song, and Y.G. Chen, Mesenchymal-epithelial interaction regulates gastrointestinal tract development in mouse embryos. Cell Rep, 2022. 40(2): p. 111053.

48. Murata, K., et al., Ascl2-Dependent Cell Dedifferentiation Drives Regeneration of Ablated Intestinal Stem Cells. Cell Stem Cell, 2020. 26(3): p. 377-390.e6.

49. Chen, L., et al., TGFB1 induces fetal reprogramming and enhances intestinal regeneration. Cell Stem Cell, 2023. 30(11): p. 1520-1537.e8.

50. Hickey, J.W., et al., Organization of the human intestine at single-cell resolution. Nature, 2023. 619(7970): p. 572–584.

51. Potten, C.S., Stem cells in gastrointestinal epithelium: numbers, characteristics and death. Philos Trans R Soc Lond B Biol Sci, 1998. 353(1370): p. 821–30.

52. Boen, C.E., et al., Patterns and Life Course Determinants of Black-White Disparities in Biological Age Acceleration: A Decomposition Analysis. Demography, 2023. 60(6): p. 1815–1841.

53. Yannatos, I., et al., Epigenetic age and socioeconomic status contribute to racial disparities in cognitive and functional aging between Black and White older Americans. medRxiv, 2023.

54. Shamsuddin, A.a.Y., GY, The large intestinal mucosa. In Whitehead R, ed. Gastrointestinal and Oesophageal Pathology, 2nd edition. New York: Churchill Livingstone. 1995. 42.

55. Nicholson, A.M., et al., Fixation and Spread of Somatic Mutations in Adult Human Colonic Epithelium. Cell Stem Cell, 2018. 22(6): p. 909-918.e8.

56. Liu, Y., et al., Comparative Molecular Analysis of Gastrointestinal Adenocarcinomas. Cancer Cell, 2018. 33(4): p. 721-735.e8.

57. Pohl, H., et al., Association between adenoma location and risk of recurrence. Gastrointest Endosc, 2016. 84(4): p. 709–16.

58. Chen, B., et al., Differential pre-malignant programs and microenvironment chart distinct paths to malignancy in human colorectal polyps. Cell, 2021. 184(26): p. 6262-6280.e26.

59. Lin, Y., et al., Normal breast tissues harbour rare populations of aneuploid epithelial cells. Nature, 2024.

60. Dart, A., Peto’s paradox put to the test. Nat Rev Cancer, 2022. 22(3): p. 129.

61. Tomasetti, C. and B. Vogelstein, Cancer etiology. Variation in cancer risk among tissues can be explained by the number of stem cell divisions. Science, 2015. 347(6217): p. 78–81.

